# Variant characterization in the intrinsically disordered human proteome

**DOI:** 10.1101/2025.06.27.661911

**Authors:** Dalmira Hubrich, Jesus Alvarado Valverde, Chop Yan Lee, Milena Djokic, Mareen Welzel, Kristina Hintz, Katja Luck

## Abstract

Variant effect prediction remains a key challenge to resolve in precision medicine. Sophisticated computational models that exploit sequence conservation and structure are increasingly successful in the characterization of missense variants in folded protein regions. However, 37% of all annotated missense variants reside in 25% of the proteome that is intrinsically disordered, lacking positional sequence conservation and stable structures. To significantly advance variant effect prediction in disordered protein regions, we combined sequence pattern searches with AlphaFold and experiments to structurally annotate 1,300 protein-protein interactions with interfaces mediated by short disordered motifs binding to folded domains in partner proteins. These interfaces were selected based on their overlap with uncertain missense variants enabling reliable prediction of deleterious effects of 1,187 uncertain variants in disordered protein regions. Extensive experimental efforts validate predicted interfaces and deleterious variant effects that were predicted as benign by AlphaMissense. This study demonstrates how critical structural information on protein interaction interfaces is for variant effect prediction especially in disordered protein regions and provides a clear avenue towards its system-wide implementation.

## Introduction

Sequencing of whole genomes, exomes, or gene panels enables identification of likely causal genetic variants, which is critical for deriving patient-specific diagnosis and the selection of tailored therapies. However, most causal genetic variants are ultra-rare in the human population and dozens of mutations arise de novo in every germline requiring very large cohort studies for the use of statistical association to discriminate benign from pathogenic mutations^1,2^. And yet, only a minority of patients suffering from rare diseases will receive a genetic diagnosis this way^2,3^. The vast majority of identified variants remain uncharacterized, hindering diagnosis, treatment, and the advancement of our mechanistic understanding of human disease^1^. Many machine learning and artificial intelligence (AI)-based methods have been developed to aid in the prediction of variant effects in protein coding sequences based on exploring sequence conservation and structural features of proteins. While these features work reasonably well to call out likely deleterious mutations in folded protein regions^4,5^, about 25% of the human proteome is estimated to be intrinsically disordered^6^. Intrinsically disordered protein regions (IDRs) are implicated in numerous critical functions in cell signaling and regulation, yet because these regions lack a defined tertiary structure, functional conservation is not expressed in precise positional conservation of amino acids leading to an overall lower accuracy of variant effect prediction in IDRs^4,5^.

Many molecular functions of IDRs are mediated via binding to other biomolecules such as proteins^6^. A well-studied mode of binding involving IDRs are short linear motifs (SLiMs, hereafter referred to as motifs) that are usually 4-10 residues long and may adopt a secondary structure upon binding to folded domains in partner proteins^7^. These domain-motif interfaces (DMIs) function in the scaffolding of signaling complexes, in mediating substrate recognition by enzymes such as E3 ligases, kinases, or phosphatases, encode sites for subcellular localization or post-translational modifications^8^. Disruption of functional motifs by mutations has been associated with cancer and Mendelian-like genetic diseases^7,9,10^. Likewise, genetic variation in IDRs can also lead to novel motifs and gain-of-function effects, as described for de novo dileucine motifs in cytosolic IDRs of transmembrane proteins^11^. In general, pathogenic mutations in IDRs were found to be enriched in motifs compared to neutral mutations^12,13^ and in a systematic screen dozens of pathogenic mutations in motifs were found to be disruptive to protein binding supporting the relevance of motifs for human disease^14^.

DMIs are estimated to be highly prevalent in the human protein interactome^15^. While we have seen major advances in the systematic mapping of human protein-protein interactions (PPIs)^16,17^, for more than 90% of those we lack structural information^16,18^, which we consider critical for more accurate variant effect prediction in IDRs. AI-based structure prediction tools such as AlphaFold (AF) enabled major advances in the structural annotation of protein interactomes^19^, yet perform much more poorly in the prediction of protein interaction interfaces that involve IDRs such as motifs, especially when using full length proteins or longer protein fragments for structure prediction^20,21^. To overcome this limitation, we had previously developed and applied an in silico protein fragmentation approach where ordered protein fragments are paired with short disordered protein fragments from an interacting pair of proteins for structural modelling by AF. While this approach coupled with experimental validation resulted in the identification of novel DMIs, it is computationally too expensive to be done on thousands of PPIs^20^.

Here, we developed a simple machine learning approach to prioritize protein fragments for structural modelling of PPI interfaces that likely contain the domain and motif mediating the interaction. We applied this DMI predictor onto more than 50,000 human PPIs followed by structural modelling of thousands of predicted DMIs that overlapped with uncertain or pathogenic variants. Experimental validation confirmed novel DMIs and deleterious effects of pathogenic and uncertain variants in motifs and domains. Importantly, experimentally proven deleterious variants in motifs are predicted to be benign by AlphaMissense, a state-of-the-art tool for variant effect prediction^22^. Using confident structural models of DMIs we predict for more than 1,000 uncertain variants in motifs to be disruptive to protein function representing a major advance and path forward towards more accurate variant effect prediction in the disordered human proteome.

## Results

### Identification of missense variants in motif-mediated protein interfaces

According to ClinVar^23^, 83% (822,942) of all annotated missense variants (in the following referred to as variants) are of uncertain significance (Fig. 1a). Of those, 37% (361,081) reside in 25% of the proteome that is intrinsically disordered (Fig. 1b, Extended Data Fig. 1a) underlining our interest in the development of approaches to better predict the functional impact of missense variation in IDRs. To this end, we developed a Random Forest Model (referred to as DMI predictor) that predicts with high accuracy (AUC=0.94, AP=0.93) likely interacting domain and motif regions for a given pair of interacting proteins (Fig. 1c-d, Extended Data Fig. 1b)^24^. Predictions are restricted to occurrences of 291 known DMI types that were manually curated from the ELM resource^25^ (Table S1). To predict disordered protein regions that likely mediate protein binding, we applied our DMI predictor to more than 50,000 published binary and likely direct human PPIs that resulted from yeast two-hybrid screens (the HuRI dataset)^16^, resulting in 12,957 predicted DMIs among 3,188 PPIs using a DMI predictor cutoff of 0.7, corresponding to an estimated sensitivity of 66% and specificity of 97% (Fig. 1e, Extended Data Fig. 1c). DMI predictions distribute well among six major functional motif categories and cover 64.5% of all annotated human DMI types (Fig. 1e, Extended Data Fig. 1d-e). 31 predicted DMIs are previously known according to annotations in ELM (Extended Data Fig. 1f).

**Figure 1.**
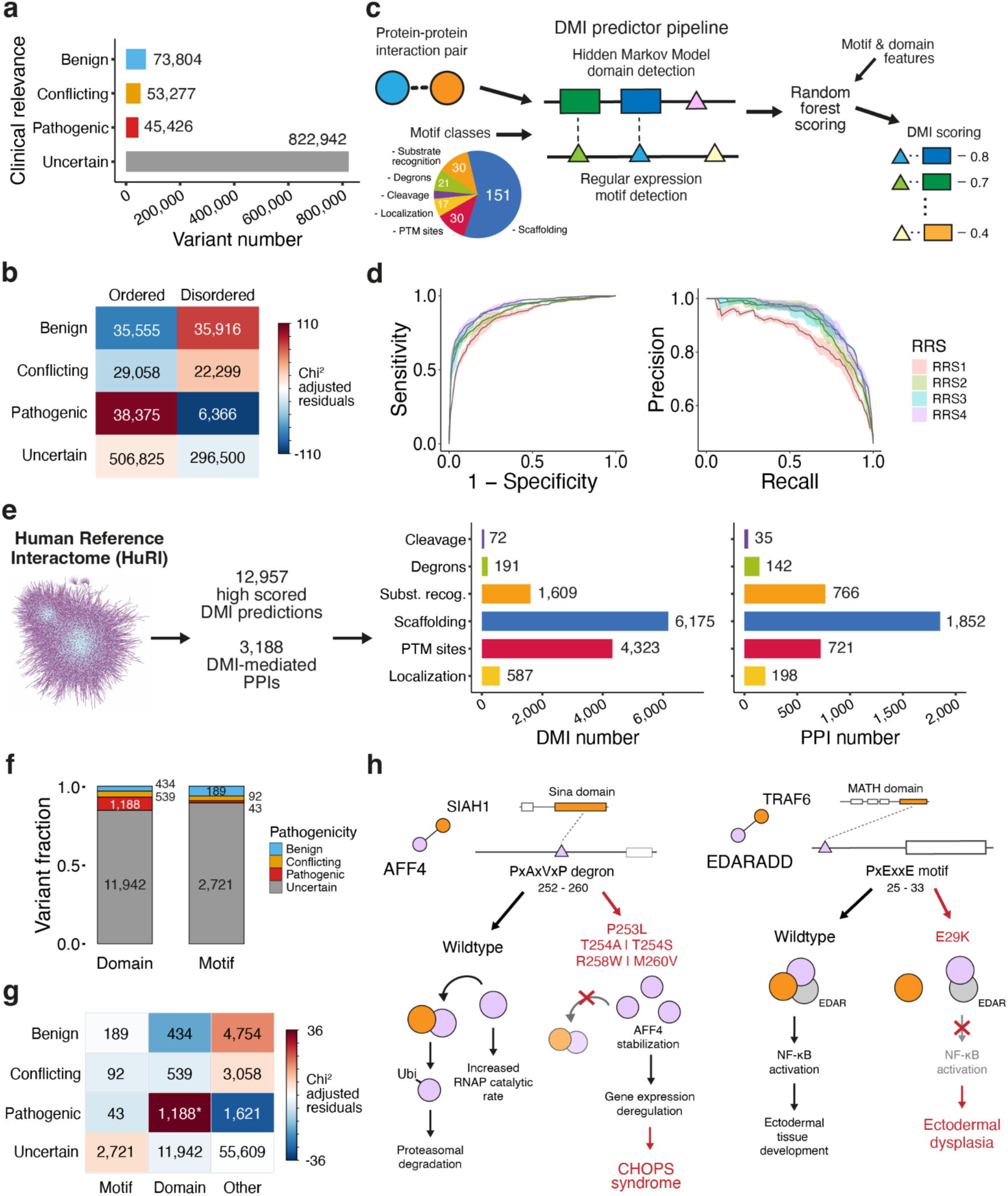
Prediction of DMIs and overlap with genetic variants. **a.** Total number of missense variants annotated in ClinVar by significance category. **b.** Chi-square frequency test residuals and count matrix for variants between ordered and disordered protein regions. Chi^2^=16,199, df=3, p-value<2.2e-16. **c.** Schematic illustrating the DMI predictor workflow. **d**. ROC and PR curves evaluating the performance of the DMI predictor using four different versions of random reference sets (RRS). Performance is averaged across three evaluations using different RRS version assemblies. **e**. Total number of predicted DMIs and corresponding number of PPIs from HuRI^16^ with at least one predicted DMI, and split by motif category according to ELM^25^. **f**. Proportion and number of variants overlapping with domains and motifs from predicted DMIs. **g.** Chi-square frequency test residuals and count matrix by ClinVar significances and predicted motifs, motif-binding domains, and other protein regions. Chi^2^=1,611, df=6, p-value<2.2e-16. *highlights a statistically significant difference between observed and expected counts (Bonferroni-adjusted p<0.05). **h.** Schematic of examples of PPIs with predicted and known DMIs that overlap with pathogenic variants in the motif and downstream consequences of PPI disruption.

Using the start and end positions of domain and motif occurrences within the predicted DMIs we find that 10,238 predicted DMIs among 2,624 PPIs overlapped with at least one variant of which the large majority involves overlap with uncertain variants (Fig. 1f, Extended Data Fig. 1g). We observe a mild overrepresentation of uncertain variants in predicted motifs while known pathogenic variants are, as expected, significantly enriched among predicted domains (Fig. 1g). 43 pathogenic variants overlapped with motifs among our predicted DMIs corresponding to different functional classes (Extended Data Fig. 1h). Closer inspection of these pathogenic variants revealed known and putative disease mechanisms involving mutated motifs (Fig. 1h). For instance, five pathogenic variants overlapped with a predicted and previously known degron motif in AFF4 that is recognised by the E3 ligase SIAH1^26^. Izumi et al showed that T254A, T254S, and R258W stabilised AFF4 protein levels, leading to increased gene expression levels of the AFF4 target genes *MYC* and *JUN*^27^. Overall, their study associated these and other mutations in AFF4 with a novel severe neurodevelopmental disorder called CHOPS (Fig. 1h).

Another pathogenic mutation E29K overlaps with a predicted and known TRAF6-binding motif in the protein EDARADD. EDARADD forms a complex with the receptor protein EDAR via a DEATH-DEATH domain interface^28^ and with TRAF6 via the motif in EDARADD that is recognised by the MATH domain in TRAF6^29^. This complex is crucial to activate NFKB signalling, as EDAR is unable to interact with TRAFs by itself^30^. This interaction directs the development of ectodermal tissues such as hair, teeth, and sweat glands^31^. Mutations in the ligand EDA, EDAR, or EDARADD are associated with hypohidrotic ectodermal dysplasia (HED). While functional studies have so far associated mutations in the EDARADD DEATH domain with HED^32,33^, the functionally unstudied variant E29K, which disrupts a highly conserved Glutamate in the TRAF6-binding motif pattern, likely disrupts NF-kB activation alike via perturbation of the interaction with TRAF6 (Fig. 1h).

Y705 in STAT3 is phosphorylated in response to cytokine or growth factor stimulation. This phosphorylation creates an SH2 domain-binding site, which is recognised by the SH2 domain of another STAT3 molecule, leading to STAT3 homodimerization, STAT3 cytoplasmic to nuclear relocation, and activation of target gene expression^34^. Y705F was shown to disrupt STAT3 homodimerization and, in a dominant negative way, impair the proper function of wildtype STAT3 in cells^35^. Mutations in the DNA-binding domain and SH2 domain of STAT3 have been associated with Hyper-IgE syndrome 1 as well as autoimmune disease^36^. Various (likely) pathogenic mutations are located at or close to position 705, such as Y705C and the uncertain variant Y705H, which we speculate are very likely to disrupt STAT3 signaling similar to Y705F. These examples illustrate known and putative cases where disruption of motifs due to missense mutations in IDRs cause different types of genetic diseases, further supporting our approach in the use of DMIs to predict deleterious effects of uncertain variants in IDRs.

### Experimental validation of predicted DMIs

To experimentally validate predicted DMIs that are likely of disease relevance, we prioritized predicted DMIs that overlapped with uncertain variants, preferentially in the motif, and where the corresponding domain-containing protein had multiple partners with predicted or known motifs or alternatively unknown modes of protein binding (Fig. 2a). We used AlphaFold to predict for all selected PPIs a structure of the complex (Table S2, Dataset S1). In cases of known or predicted DMIs, we used the corresponding protein fragments for structural modelling. In cases of unknown modes of protein binding, we employed our previously published *in silico* fragmentation and AF screening strategy^20^. Resulting models were evaluated based on their model confidence and manual inspection. Likely accurate models were used to design point mutations of interface residues aimed at disrupting interactions without interfering with protein stability or folding. We complemented point mutations with deletions of predicted or known motifs. Wildtype and mutant proteins were assayed for binding using bioluminescence resonance energy transfer (BRET) titration experiments^37^ (Table S3, Dataset S2). Differences in binding between wildtype and mutant were quantified by differences in estimated BRET50 or maxBRET (Fig. 2b, Table S4,S5).

**Figure 2.**
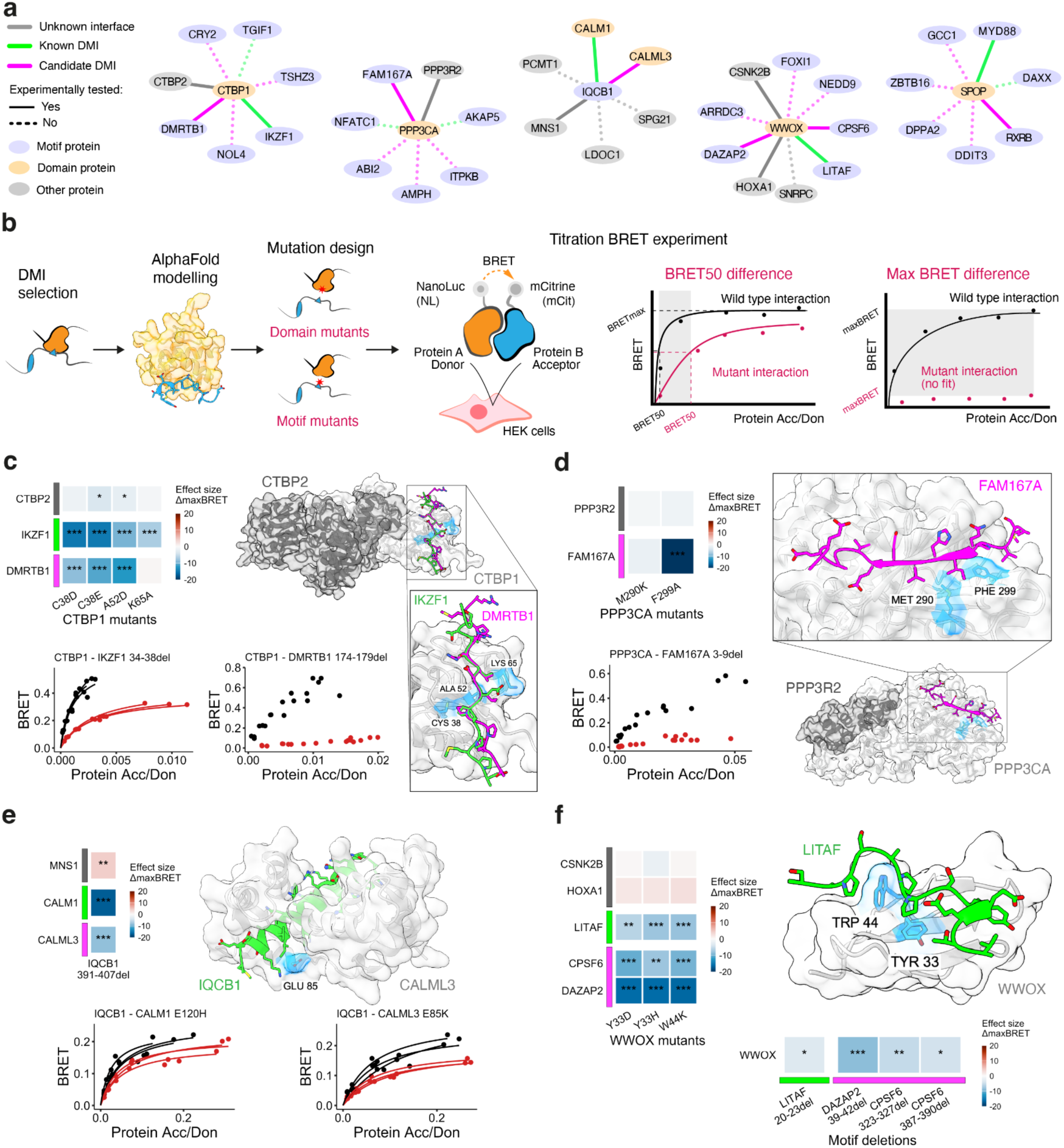
Experimental validation of predicted DMIs. **a.** Proteins and interactions selected for experimental validation of predicted DMIs. **b.** Schematic illustrating experimental approach to validate predicted DMIs. **c**. Heatmap of effect sizes based on T-test of maxBRET measurements obtained for wildtype versus mutant CTBP1-partner interactions validating the motif-binding pocket on CTBP1. *** p-value < 0.001, ** p-value < 0.01, * p-value < 0.05, three technical replicates each. Effect size values capped at 20 and -20. Color bars on y-axis: grey=other interface, green=known DMI, magenta=predicted DMI. BRET titration curves of CTBP1 versus wildtype (black) and mutant IKZF1 (red, left, ΔBRET50 effect size=-7.06, p-value=0.0021 based on T-test) and DMRTB1 (red, right, ΔmaxBRET effect size=-9.32, p-value=8.6e-05 based on T-test) for which the known and predicted motif was deleted, respectively. Data is shown for three technical replicates. Superimposed structural models of the CTBP domain of CTBP1 (14-365) with CTBP2 (21-365) and PxDLS motifs in IKZF1 (29-43) and DMRTB1 (169-183). Domain residues mutated to validate the motif-binding pocket are highlighted in blue. **d.** Same as in **c** but for experimental validation of predicted DMI between PPP3CA and FAM167A, ΔmaxBRET effect size=-20.9, p-value=6.07e-09 for PPP3CA vs. FAM167 wildtype and motif deletion. Structural model of the Metallophos domain of PPP3CA (13-485) with PPP3R2 (12-170) and the motif in FAM167A (1-16). **e.** Same as in **c** but for predicted DMI between CALML3 and IQCB1, ΔBRET50 and ΔmaxBRET not significant for IQCB1-CALM1 wildtype vs. E120H; ΔmaxBRET effect size=-6.86, p-value=2.36e-03, ΔBRET50 not significant for IQCB1-CALML3 wildtype vs. E85K. Structural model of the EF-hand domain of CALM3 (1-149) with the motif in IQCB1 (374-424). **f.** Same as in **c** but for predicted DMIs involving WWOX and CPSF6 as well as DAZAP2. Structural model of the WW domain of WWOX (16-57) with the motif in LITAF (20-23).

We confirmed a known and novel PxDLS motif in partner proteins of the transcriptional co-repressor CTBP1. CTBP1 binds motifs with its CtBP domain, mediating recruitment to DNA^38,39^. Via BRET titrations, we found that designed point mutations in the motif-binding pocket of the CTBP1 CtBP domain significantly reduced binding to the known and predicted motif partner protein IKZF1 and DMRTB1, respectively, while only mildly affecting binding to CTBP2 (Fig. 2c, Extended Data Fig. 2a). CTBP2 is known and modelled to bind to CTBP1 via a heterodimerization interface on the CtBP domains of both proteins^40^, which does not overlap with the motif-binding pocket supporting that the mutations in the motif-binding pocket specifically disrupted interaction with motif partner proteins (Fig. 2c). Interestingly, motif pocket mutation K54A only disrupted binding to IKZF1 but not DMRTB1. AF-generated structural models suggest that K54 exclusively mediates a salt bridge with E35 in the IKZF1 motif, illustrating the sensitivity of the BRET assay and accuracy of AF models. Furthermore, we observed that deletion of the known and predicted PxDLS motif in IKZF1 and DMRTB1, respectively, significantly reduced binding to CTBP1 (Fig. 2c), overall validating the predicted DMI between CTBP1 and DMRTB1 and associating the uncharacterized protein DMRTB1 with CTBP1-mediated transcription repression activity. Similar results were obtained for the predicted calcineurin (PPP3CA) docking motif in FAM167A (Fig. 2d, Extended Data Fig. 2a), and the novel predicted DMI involving the known IQ motif in IQCB1 and the calmodulin-like protein CALML3 (Fig. 2e, Extended Data Fig. 2a). We furthermore validated novel DMIs between WWOX and WW domain-binding motifs in DAZAP2 and CPSF6 (Fig. 2f, Extended Data Fig. 2a).

The E3 ligase SPOP recognizes protein substrates via degrons that are annotated in ELM to correspond to [AVP].[ST][ST][ST] sequence patterns where residues within brackets indicate possible residues at this position and a dot corresponds to a position accepting any amino acid^41^. We predicted two SPOP degrons in the partner protein RXRB, however, failed to predict the known degron of sequence VDSS in MYD88 because it lacks the third presumably required Serine or Threonine (Extended Data Fig. 2b)^42^. We identified an additional potential SPOP degron in MYD88 close to the N-terminus, which, however, is poorly conserved and did not meet our DMI predictor cutoff (Extended Data Fig. 2b). Motif pocket mutations as well as motif deletions validated the predicted motifs in RXRB and the known motif in MYD88 but failed to confirm the true negative N-terminal degron in MYD88 (Extended Data Fig. 2c). We furthermore observed that joint deletion of both motifs in RXRB only resulted in mild disruption of binding to SPOP. Further inspection of the RXRB sequence identified in total four potential SPOP degrons of which two, like MYD88, only partially matched our motif pattern (Extended Data Fig. 2d). These results illustrate on one hand, some limitations in the use of restrictive regular expressions for finding functional motif matches, but on the other hand, also demonstrate specificity in our predictions in ranking likely false motifs more lowly. Overall, our successful experimental validations of multiple predicted and known DMIs now form the basis for subsequent identification of uncertain variants in motifs that likely disrupt protein function.

### Experimental assessment of variant effects

We identified an uncertain variant R178H within the novel CTBP1-binding motif in DMRTB1 as well as an uncertain variant V8M within the novel calcineurin-binding motif in FAM167A, which both strongly perturbed binding to their partner proteins in BRET titration experiments (Fig. 3a-b, Extended Data Fig. 2e). For both uncertain variants no associated conditions have been annotated. Contrary to these variants in motif core positions, we found that three uncertain variants (M31V, S41L, T39S) in the flanking regions of the CTBP1-binding motif in IKZF1 had much milder effects on CTBP1 binding (Fig. 3a, Extended Data Fig. 2e). Interestingly, two uncertain variants overlapping with the IQ motif in IQCB1, R404G found in a patient with Nephronophthisis and N406Y, only significantly affected binding to CALML3 but not CALM1 (Fig. 3c, Extended Data Fig. 2e). Inspection of the structural models of these DMIs did not reveal any obvious properties explaining these different effects. We also identified two uncertain variants in the motif-binding pocket of CALM1, which we found disruptive to IQCB1 binding (Fig. 3d, Extended Data Fig. 2e). R87H was reported in a patient with a cardiovascular phenotype according to ClinVar and D94H was reported in patients with Catecholaminergic polymorphic ventricular tachycardia 4 and Long QT syndrome 14^23^. The uncertain variant A89D in the motif-binding pocket of CALML3 that also significantly weakened binding to IQCB1 (Fig. 3d, Extended Data Fig. 2e), is reported without disease annotations.

**Figure 3.**
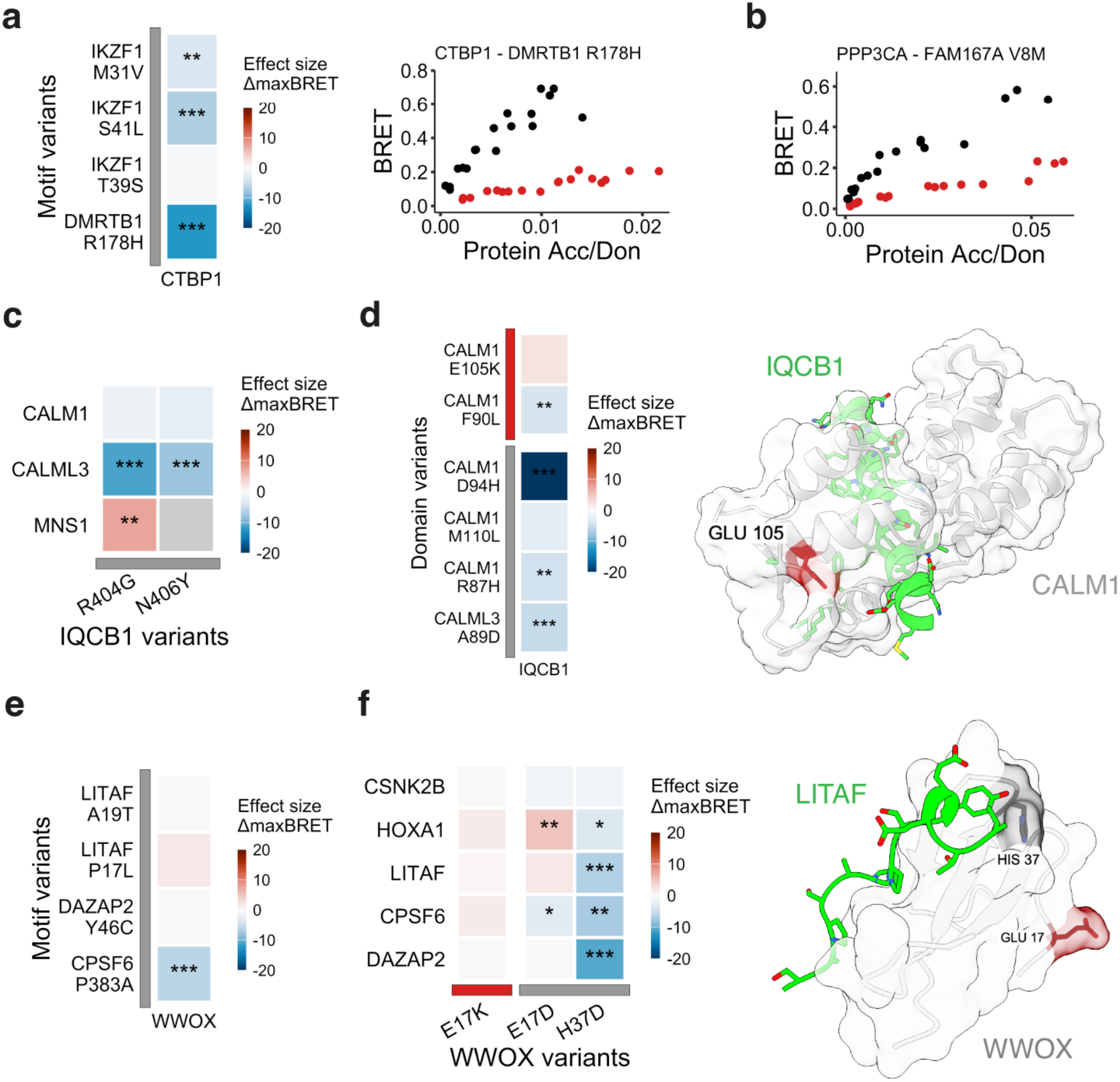
Experimental testing of variants in DMIs. **a.** Heatmap of effect sizes based on T-test of maxBRET measurements obtained for wildtype versus uncertain variants in CTBP1 partner proteins assayed for binding against wildtype CTBP1. *** p-value < 0.001, ** p-value < 0.01, * p-value < 0.05, three technical replicates each. Effect size values capped at 20 and -20. BRET titration curves of CTBP1-DMRTB1 wildtype (black) interaction vs. DMRTB1 R178H (uncertain variant, red, maxBRET T-test effect size=-12.19, p-value=5.7e-06, 3 technical replicates). **b.** BRET titration curves of PPP3CA-FAM167A wildtype (black) interaction vs. FAM167A V8M (uncertain variant, red, maxBRET T-test effect size -24.23, p-value 3.2e-07, 3 technical replicates). **c.** Same as in **a** but for uncertain variants in IQCB1. **d.** Same as in **a** but for pathogenic (red) and uncertain variants (grey) in IQCB1-partner proteins. Structural model of the EF-hand domain of CALM1 (1-149) with the motif in IQCB1 (374-424). Position of pathogenic variant E105K is highlighted in red. **e.** Same as in **a** but for uncertain variants in WWOX-partner proteins. **f.** Same as in **a** but for pathogenic (red) and uncertain variants (grey) in WWOX. Structural model of the WW domain of WWOX (16-57) with the motif in LITAF (20-23). Domain positions H37 and E17 are highlighted according to the significance of annotated variants at these positions.

We further tested the effect of two pathogenic mutations in the motif-binding pocket of CALM1. E105K received its pathogenic annotation based on computational assessments^23^. We observed that this mutation slightly enhanced binding to ICQB1 likely because the Glu to Lys mutation removes repulsive negative charges and enables salt bridges with Glu residues on the IQ motif in IQCB1 (Fig. 3d, Extended Data Fig. 2e). The pathogenic mutation F90L identified in patients with idiopathic ventricular fibrillation and early sudden cardiac death weakened binding to IQCB1 (Fig. 3d, Extended Data Fig. 2e) and based on previous studies was found to decrease Ca2+ binding and interaction with RyR2^43^. Unfortunately, no uncertain variants overlapped with DMIs involving the Cullin E3 ligase SPOP. However, the pathogenic mutation G132V in the motif-binding domain of SPOP that was identified in a patient with macrocephaly, cardiovascular abnormalities and intellectual disability and reported to lead to diminished partner binding^44^, also significantly reduced binding to RXRB and MYD88 (Extended Data Fig. 2f).

No pathogenic or uncertain variants overlapped with known or novel WW-binding motifs in LITAF, CPSF6, or DAZAP2. Uncertain variants in the flanking regions of these motifs did not or only very mildly affect binding to WWOX (Fig. 3e, Extended Data Fig. 2e). Interestingly, LITAF has two known PPSY motifs (Extended Data Fig. 3a), of which the second motif (58-61) was reported previously not to contribute to WWOX binding^45^. Accordingly, mutations aimed at disrupting the second motif or uncertain variants overlapping with this second motif mostly did not reduce binding to WWOX (Extended Data Fig. 3b). We furthermore identified one pathogenic and two uncertain variants in the motif-binding pocket of the first WW domain of WWOX (Fig. 3f). E17K is annotated as pathogenic based on computational assessments while E17D is uncertain. E17 is on the surface of the WW domain and away from the motif-binding pocket. Accordingly, we found that both variants did not or only very mildly interfere with binding to any of the five tested WWOX partner proteins (Fig. 3f). Conversely, the uncertain variant H37D in the motif-binding pocket specifically disrupted binding to the three PPxY motif-containing partner proteins but not to CSNK2B and only very mildly to HOXA1, which both do not contain a PPxY motif (Fig. 3f). All three mutations had been identified in patients displaying developmental and epileptic encephalopathy or/and autosomal recessive spinocerebellar ataxia 12^23^ but our data suggest that only H37D might be deleterious to WWOX function.

Overall, we observed that measured effects of variants on protein binding seemed to correlate well with predictions based on the structural models obtained for each DMI. To quantify this observation and generate a predictive model of variant effects based on structural models of DMIs, we first determined motif and domain residues at the interface based on changes in the relative surface accessibility (ΔRSA) between bound and unbound forms of the motifs and domains. Interestingly, we observed that tested variants in the core and flanking motif residues both displayed equal ΔRSA distributions suggesting that both are equally important for domain binding (Fig. 4a). However, based on our BRET measurements, we found that mutations in flanking motif positions were less disruptive to partner binding (Fig. 4b). In line with previous reports^20^, we conclude that AF places flanking residues of core binding regions close to partner residues while being less confident in this prediction (Fig. 4c). We therefore decided to not only consider structural aspects but also model accuracy in our classification of motif residues into interface vs not interface and flanking positions. Using ΔRSA and RSA values we further classified domain residues into interface and otherwise surface and buried positions (Fig. 4d). Using this residue classification, we find that domain and motif interface as well as buried domain variants (apart from one gain of function mutation) are more disruptive to protein binding based on BRET measurements compared to domain surface and motif non-interface variants (Fig. 4e).

**Figure 4.**
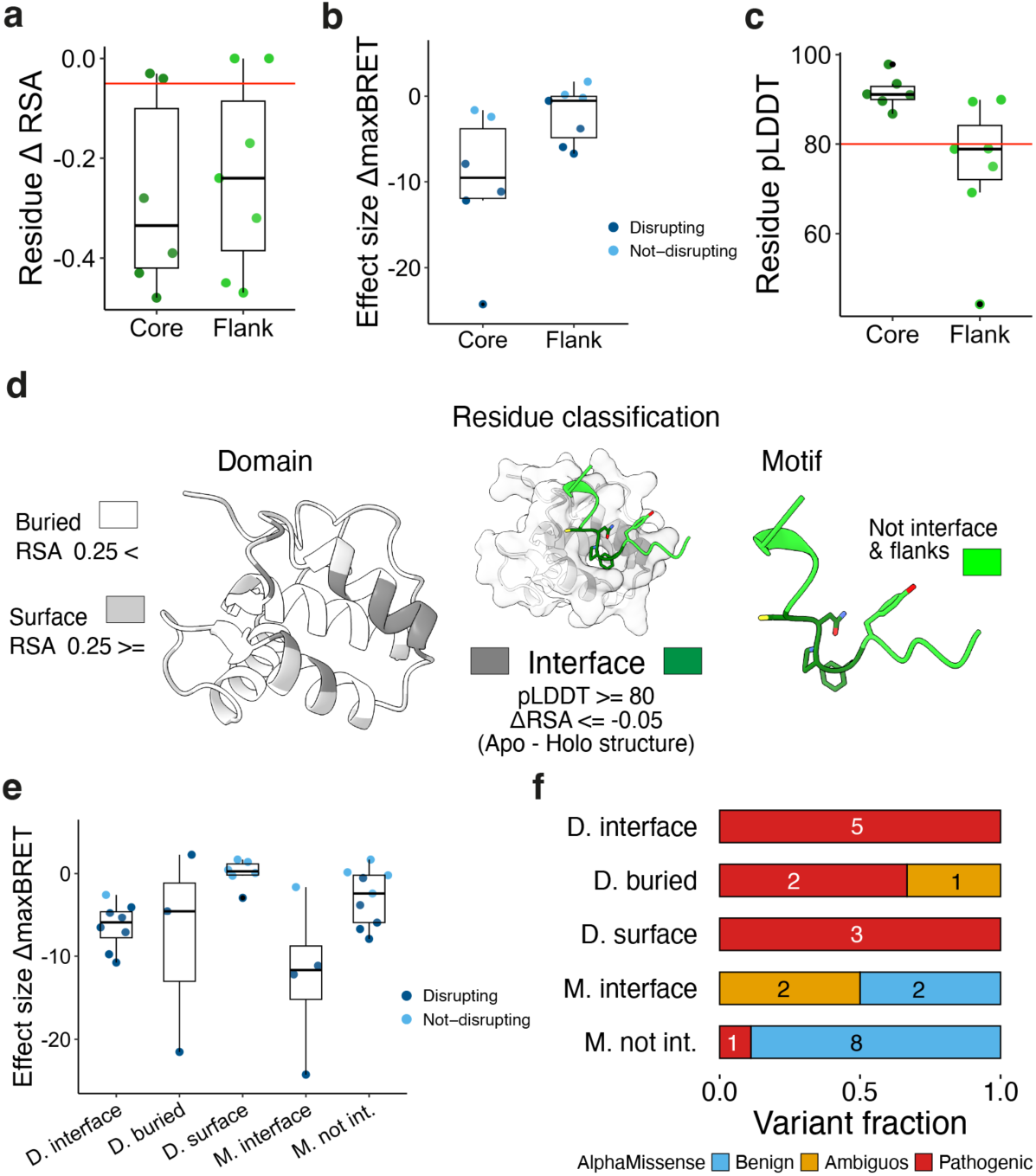
Structure-based residue classification. **a**. ΔRSA scores of core and flank motif positions that carry experimentally tested variants (red line at ΔRSA=-0.05 indicates threshold below which residue positions are considered part of the interface). **b.** Effect sizes based on T-test of ΔmaxBRET measurements between wildtype and variants at motif core and flank positions. A variant is labelled as disruptive if ΔmaxBRET was significant. **c**. pLDDT scores of residue positions in motif core and flanks that carry experimentally tested variants (red line at pLDDT=80 indicates threshold above which residues are considered to be confidently modelled). **d.** Schematic illustrating classification of domain and motif residue positions based on DMI structural models. **e.** Effect sizes based on T-test of ΔmaxBRET measurements between wildtype and variants split by classification of residue position in DMI structural models; D.=domain, M.=motif, not int.=not interface. **f.** Fraction and number of experimentally tested variants split by residue classification according to DMI structural models and further grouped based on AlphaMissense predictions. Abbreviations as in **e**. AlphaMissense (AM) ranks top among other tools for variant effect prediction^4,22^. We compared AM predictions with our structure-based classification for tested variants and found that AM tends to predict mutations within domains as likely pathogenic regardless of their position, i.e. surface vs. buried. Conversely, variants in motifs are generally predicted to be benign regardless if they correspond to motif interface positions or not (Fig. 4f). This suggests that structural information of PPIs for the prediction of variant effects, especially in disordered protein regions, is critical to achieve more accurate pathogenicity predictions.

### Variant effect prediction in the disordered proteome

Encouraged from these results we generated structural models for 7,341 of our predicted DMIs that overlapped with at least one pathogenic or uncertain variant (Table S6). Based on previously published benchmarking efforts^20^ we applied a model confidence cutoff of 0.6 to retain 3,450 DMI structural models corresponding to 1,300 PPIs (Fig. 5a). Successfully modelled DMIs correspond to different functional classes (Extended Data Fig. 3c) and confirm earlier observations that flanking residues of motifs display a sharp decrease in pLDDT compared to a much less pronounced increase in ΔRSA (Extended Data Fig. 3d). Using these structural models we classified all domain residues into either buried, surface, or interface as well as motif residues into interface or non-interface using the strategy described earlier (Fig. 4d). Overlapping this structural classification of residue positions with variant annotations from ClinVar we can predict interface disruption as a pathogenic mechanism for 193 pathogenic variants overlapping with domain interface positions while another 1,826 pathogenic variants overlapping with domain buried positions likely lead to fold destabilization (Fig. 5b). Most importantly, 52.7% (11,906) out of all uncertain variants that overlapped with any domain or motif position are predicted to be deleterious to protein function based on their location at buried or interface positions (Fig. 5b, Table S7, Dataset S3). Out of those, 1,187 uncertain variants fall into motif interface positions. Of note, while AM predicts most variants at domain interface positions to be pathogenic, it evaluates most variants at motif interface positions to be benign (Fig. 5c).

**Figure 5.**
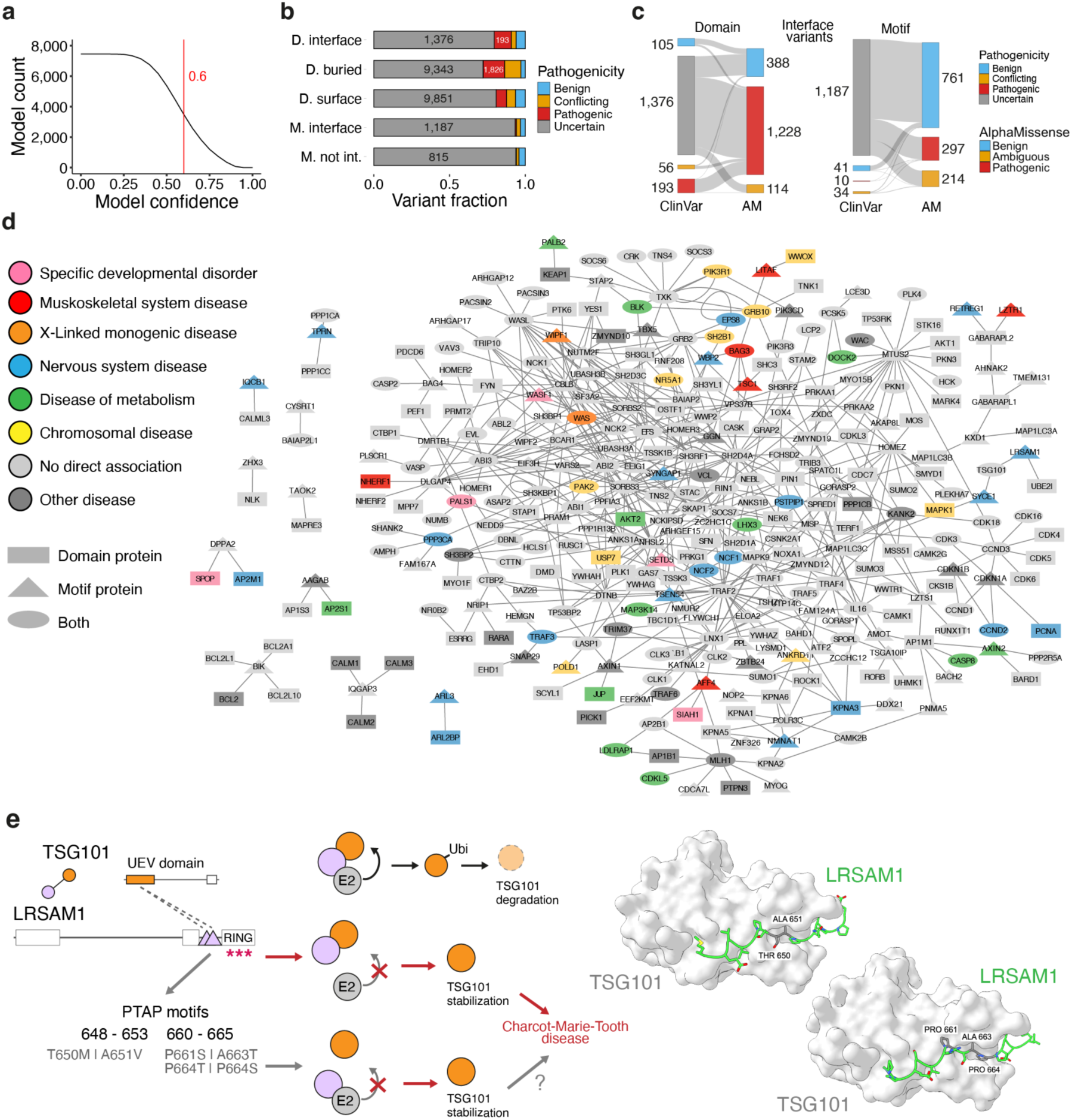
Proteome-wide prediction of variant effects in IDRs. **a.** Number of structurally modelled DMIs for increasing model confidence cutoffs. The red line indicates the chosen model confidence cutoff of 0.6 corresponding to 3,450 confidently predicted models. **b.** Fraction and count of variants by ClinVar significance and residue structural classification. Variants at interface or buried positions are likely deleterious. **c.** Sankey plot of variants predicted to be at domain or motif interface positions split by ClinVar significance and AlphaMissense prediction. **d.** Network of PPIs with DMIs with at least one uncertain variant in a predicted motif interface position. Disease annotations extracted from OrphaNet for rare diseases. **e.** Schematic of LRSAM1-TSG101 interaction, protein architecture, and molecular function. Known pathogenic mutations in the LRSAM1 RING domain as well as uncertain variants in predicted interfacial motif positions likely stabilize TSG101 protein levels leading to disease. Structural models of TSG101 UEV domain (1-145) and LRSAM1 PTAP motifs (643-657, 655-669) are shown with predicted motif interface residues in grey carrying uncertain variants.

Interacting proteins predicted to be mediated by a structurally modelled DMI with an uncertain variant at a motif interface position carry various disease annotations and form the basis for further downstream investigations of potential molecular disease mechanisms (Fig. 5d). For instance, within this network there is the interaction between the E3 ligase LRSAM1 and TSG101 that is mediated by two known occurrences of the PTAP motif in LRSAM1 (Extended Data Fig. 3e) binding to the UEV domain in TSG101. As a consequence, the LRSAM1 ubiquitinates TSG101 leading to its proteasomal degradation^46^ (Fig. 5e). Dysregulation of TSG101 protein levels via non-functional LRSAM1 has been implicated in Charcot-Marie-Tooth (CMT) disease^47^. However, so far only mutations disrupting the function of the C-terminal RING domain in LRSAM1 have been linked to CMT^47,48^. We find numerous uncertain variants identified in patients with CMT^23^ to overlap with predicted interface positions in both motifs suggesting that not only perturbation of the interaction between the E3 and E2 enzymes because of RING domain loss deregulates TSG101 levels but equally alike perturbation of the direct interaction between LRSAM1 and TSG101. Of note, AM evaluated these motif variants as benign.

## Discussion

Genotype-to-phenotype relationships are immensely complex even at the molecular level when we restrict ourselves to predicting the effect of a coding missense mutation on protein function. Proteins are inherently complex molecules where individual residues function in folding, binding, catalytic reactions, or contribute to the overall architecture of the protein, i.e. by placing functional regions within the right distance or orientation to each other. Some of these functions, such as folding, can be predicted reasonably well from primary sequence conservation or from monomeric structural models^49^. However, in the absence of structural models of protein complexes our work strongly suggests that even sophisticated neural network models fail to accurately predict deleterious effects of mutations in PPI interfaces. This limitation is particularly problematic for the characterization of genetic variation in disordered protein regions, which most often function by binding partner molecules and where protein folding plays little importance^6^. To overcome this limitation, we developed a simple yet powerful computational approach that guides structural modelling of disordered PPI interfaces by AF using sequence pattern searches and other protein features combined in a Random Forest model. The 3,450 confidently structurally modelled DMIs by this approach represent ample opportunities to study molecular mechanisms of cell regulation with potential disease relevance. The scope of the DMI predictor can be improved in future studies by including more curated DMI types, i.e. from the MoMap resource (slim.icr.ac.uk/momap/), by expanding to other PPI resources, and by incorporating more flexible motif definitions, i.e. by using position weight matrices. The success of this approach also underlines the importance of efforts cataloging PPI interface types^25,50^.

Experimental tests confirmed six new DMIs, deleteriousness of eleven uncertain variants, and a likely benign effect of one pathogenic mutation. For most of these variants we provide first experimental data to assess their effects. Our work further demonstrated that structural models of protein complexes generated by AF are accurate enough to design point mutations at PPI interfaces that are disruptive to binding but not folding. Consequently, we showed that these structural models can be used to predict likely deleterious effects of mutations based on being in buried or PPI interface positions. Of note, this approach enabled prediction of deleterious effects for hundreds of uncertain variants in disordered protein regions. These predictions can be further improved in their accuracy by considering the type of amino acid change and not just the mutated position, i.e. by incorporating simple energetic molecular dynamics-like computations^51^. Our experimental data highlighted interaction-specific, so-called edgetic, effects of mutations^52^. Along with molecular function predictions derived from predicted DMIs, specific interface disruption provides strong hypotheses for molecular mechanisms driving observed phenotypes, a prerequisite (in the absence of statistical association) to further discern if a deleterious variant is also disease causing.

Assaying for interaction perturbation of variants in IDRs using full length proteins also highlighted the importance of multivalency, referring to multiple contact sites between two proteins, which are often formed by motifs^53,7^. Multivalent interfaces can dampen effects of variants on PPIs, which would otherwise show as disruptive when assaying a single interface using protein fragments. Future work should aim for predictive models that incorporate multivalency in variant effect predictions, especially if they occur in IDRs, and experimental validations should ideally be conducted using full length proteins, as done in this study, to confirm multivalency. Despite multiple challenges ahead, the time is ripe to systematically incorporate structural models of protein complexes in variant effect prediction bringing us a substantial leap forward in closing the genotype-phenotype gap.

## Data and Code Availability

DMI predictor code is made available on github (https://github.com/KatjaLuckLab/DMI_predictor) as well as code to process and analyze ClinVar, AlphaMissense data, and BRET data as well as AF models (https://github.com/KatjaLuckLab/DMI_ClinVar_Manuscript).

## Acknowledgements

We thank all members of the Luck lab for helpful discussions throughout the study. We thank Joelle Strom for help with BRET data analysis, Erich Wanker for discussions on BRET data analysis, and Norman Davey and Izabella Krystkowiak for help computing conservation scores for SLiM matches. We thank the protein production, media and IT core facilities from IMB. The GPU clusters on which the AlphaFold predictions were performed were funded by the Ministry of Science and Health (MWG), Rhineland Palatinate (funding ID: TB-Nr.:3658/19) and the Deutsche Forschungsgemeinschaft (DFG, German Research Foundation) – Project-ID 393547839. The research performed in this study was funded by the Deutsche Forschungsgemeinschaft (DFG, German Research Foundation) – Project-ID 449991970 and 464588647 awarded to K.L.

## Author contributions

K.L. designed the study, J.A.V, C.Y.L., M.D., K.H. performed computational work, D.H., C.Y.L., M.W. performed experiments, D.H., K.L., C.Y.L. performed experimental data analysis, K.L., J.A.V., D.H., C.Y.L., M.D., M.W. wrote the manuscript, K.L. provided the funding.

## Methods

### Processing of ClinVar missense variants

We obtained ClinVar data from the November 2023 release (https://ftp.ncbi.nlm.nih.gov/pub/clinvar/tab_delimited/archive/2023/variant_summary_2023-11.txt.gz)^23^, retaining only GRCh38 genome assembly mapped variants with valid protein-level HGVS annotations ("p."). We further selected variants occurring in canonical UniProt^54^ isoforms where the reported wild-type amino acid matched the reference sequence. Variants were then filtered by clinical significance, retaining those whose pathogenicity can be simplified as pathogenic, benign, uncertain, or conflicting. Pathogenicity categories were then simplified in the following way: “benign”, “likely benign”, “benign/likely benign” = benign and “pathogenic”, “likely pathogenic”, “pathogenic/likely pathogenic” = pathogenic. Finally, we restricted the dataset to germline missense single-nucleotide variants and excluded truncating variants ("Ter"), resulting in 996,702 variants in the final dataset.

### Classification of residue disorder

Residue disorder classification was done by using pLDDT values from all available human proteins in the AFDB^55^ (as of July 15 2024). Taking Wilson et al.^56^ as reference, a threshold value of 70 was used to define categorical groups of order (above or equal to the threshold) and disordered residues (below the threshold).

### DMI predictor development

#### Manual curation of domain-motif interface types

DMI types and their associated regular expression (REGEX) models were downloaded from the ELM DB (version 1.4, accessed on 29.09.2020)^25^. Existing annotations for interacting domains were manually checked, corrected where applicable, and missing ones added. To this end, we used information in ELM DB on known interaction partners from annotated motifs and submitted the interaction partners to SMART web server (http://smart.embl-heidelberg.de/) to check which Hidden Markov Models (HMMs) from SMART and Pfam matched with the annotated domains^57–59^. We annotated DMI types with the better matching HMM based on the domain boundaries, sensitivity and specificity of the HMM. If Pfam and SMART HMMs performed equally well for a given DMI type, then SMART HMMs were selected because overall we found them to be superior in performance. We furthermore annotated if the HMM describes a repeat from a larger domain and how many of the repeats are required to assemble the full domain that is recognizing the motif. We also curated information on cases where two different types of domains jointly recognize a motif. In total, we retained 291 fully annotated DMI types for subsequent development of the DMI predictor.

#### Positive reference DMI set curation

A list of 20,396 reviewed human protein sequences was downloaded from SwissProt (16.02.2021)^54^. A list of experimentally validated DMI instances was downloaded from the ELM database (accessed on 18.02.2021)^25^, and the list served as the basis for the curation of the positive reference set (PRS). The list was filtered for interactions involving only human proteins from SwissProt. The DMI instances in the resulting list were then subject to further processing by checking the annotated positions of the motif and domain in the interaction pairs. This was done by running the REGEX of the motif type on the sequence of the annotated motif protein to confirm the motif’s positions in the sequence. A similar step was performed on annotated domains by running HMM search on the annotated protein using the SMART web server (http://smart.embl-heidelberg.de/) and the HMM selected during the manual curation of the DMI types^57,58^. If the reported positions of the motif did not align with the match returned from pattern search, the DMI instance would be removed. Similarly, should the reported domain not occur on the annotated protein through HMM search, the DMI instance would be removed, too. DMI instances formed by alternative isoforms were converted to their canonical isoforms and manually checked to ensure that the annotated motif sequence and match positions were still the same in canonical isoforms and the annotated domains were still detected by HMMs. The resulting list contained 662 proteins that formed 898 DMI instances from 106 DMI types, serving as the PRS for the development of the DMI predictor (Table S8).

#### Random reference DMI set curation

Random reference sets (RRS) were generated by randomly pairing proteins (20,396 reviewed human protein sequences downloaded from SwissProt on 16.02.2021) while excluding known interacting protein pairs (145,468 interactions from human proteins, downloaded from IntAct on 24.02.2021). SLiM matches and domain matches were detected in randomly paired proteins by running REGEX and HMM search on the paired proteins. Potential DMIs were then matched among the paired proteins to form DMI instances for the RRS. As the PRS was made up of a relatively small pool of proteins, different strategies to sample random protein pairs were tested to ensure that the predictor was not biased by learning some inherent properties in the PRS. To this end, random protein pairs were either sampled from the PRS protein pool or the whole human proteome to allow for the comparison of prediction performance using different RRSs. Furthermore, the DMI instances in the PRS only represented a subset of all annotated DMI types. To ensure that prediction performance was measured across many DMI types, we once uniformly sampled RRS DMI instances across all DMI types. On the other hand, prediction of DMI instances depends on the frequency of domains and REGEX matches in the proteome, and sampling DMI instances uniformly would not allow these features to play in the training of the predictor. To this end, a prespecified amount of RRS DMI instances, regardless of DMI types, was sampled randomly. To sample DMI types evenly, 5 DMI instances per DMI type were sampled when the PRS protein pool was used, and 3 DMI instances per DMI type were sampled when the whole human proteome was used. The number of randomly sampled RRS DMI instances was set at 1,000 DMI instances. This step ensured that the number of DMI instances in RRS was roughly the same as that of PRS. Four versions of RRS (sampling strategies detailed below), with triplicates of each version, were generated in total by combining the strategies of sampling proteins and DMI instances (Table S8).

RRSv1: Proteins randomly paired using the PRS protein pools and 5 DMI instances uniformly sampled across all DMI types

RRSv2: Proteins randomly paired using the PRS protein pools and 1000 DMI instances sampled regardless of DMI types

RRSv3: Proteins randomly paired using the human proteome and 3 DMI instances uniformly sampled across all DMI types

RRSv4: Proteins randomly paired using the human proteome and 1000 DMI instances sampled regardless of DMI types

#### Annotation of domain and motif features for Random Forest training Motif features annotation

Features of motif and domain matches were calculated for all DMI instances in the PRS and the RRS. For motifs, we included the REGEX model probability as provided from the ELM DB. We calculated the average IUPred short and ANCHOR score across the residues of each motif match using a local installation of IUPred^60^. We determined the fraction of motif residues overlapping with any Pfam HMM domain match to identify false motif matches in folded domains. The Pfam HMM domain matches of all human proteins were retrieved from the InterProScan webserver by running Pfam HMM scan on all human proteins (accessed on 22.02.2021) (InterPro, InterProScan, Pfam)^59,61,62^. For each motif match we computed the relative local conservation (RLC) following the Sigmotif formula from SLiMPrints^63^. For the RLC, we used four types of conservation scores: conservation calculated across orthologues from Quest for Orthologs (QFO), conservation on the metazoan level, conservation on the level of class (mammalia), and subphylum (vertebrata). We additionally added the variance of the RLC scores:

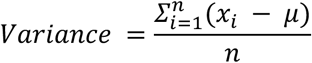

where *x*_*i*_ is the conservation score and 𝜇 the mean of conservation across the defined positions of the motif match^63^.

#### Domain features annotation

For domains, we calculated the domain frequency by protein, the domain frequency in the human proteome, and the enrichment of the domain among interaction partners. The domain frequency by protein corresponded to the fraction of proteins in the human proteome with at least one match of the domain HMM. The relative domain frequency in the human proteome corresponded to the total number of domain HMM matches divided by the total number of proteins.

#### DMI features annotation

We calculated the enrichment of the domain among the interaction partners of the motif protein compared to the background distribution of the domain in the PPI network. To this end, we downloaded and employed the HuRI and literature-curated binary PPI networks from Luck et al. (2020)^16^ and combined them to produce a PPI network consisting of 13,722 proteins and 101,219 binary interactions. Following that, 1,000 random networks were generated from the real network using the python-igraph^64^ package and degree_sequence function while maintaining the degree of the proteins observed in the real network to account for domain frequency and protein degree in the network. The enrichment of a domain among the interaction partners of a motif-containing protein was first quantified by counting the number of interaction partners having the domain. To compute the significance of the observed domain enrichment, the generated random networks were used to count the number of random interaction partners of the motif protein having the domain, thereby producing a background distribution for the domain enrichment. The p-value of the domain enrichment was then computed by calculating the fraction of random networks with greater than or equal the number of interaction partners with the domain compared to the real network. The z-score of the observed domain enrichment was also computed to quantify the extent of enrichment observed in the real network. The z-score of domain enrichment was calculated by subtracting the mean number of interaction partners with the domain in the random networks from that observed in the real network and dividing the result by the standard deviation of the number of interaction partners with the domain in the random networks. In cases where the standard deviation in the random networks is zero, the absolute difference between the number of interaction partners with the domain observed in the real network and the mean number of interaction partners with the domain in the random networks was calculated as the z-score. During the training of the DMI predictor, interacting protein pairs in the PRS were removed from the protein’s real network to avoid circularity in the calculation of domain enrichment.

Data points with missing values were removed. These removed data points were either without conservation score or PPI network information. After listwise deletion of data points with missing values, 830 DMI instances were retained in the PRS. Similar processing was applied to all versions of RRS, and the number of DMI instances retained in the triplicates of the RRS versions ranged from 870 to 985, making a fairly balanced dataset for the training of the Random Forest (RF) model.

#### Model training and testing

Each replicate of the RRS versions was combined with the PRS to train a RF model. The combined dataset was stratified by labels (known and random DMI instances) and split into a training and a test set with a ratio of 3:1. Subsequently, an RF model with 1,000 decision trees was fitted on the training set, and the fitted RF’s accuracy was then evaluated on the test set. Random seed was set to 0 for model building. Predictive performances of fitted RFs were further evaluated by means of ROC curve and PR curve. For each RRS version, ROC and PR curves were averaged across the triplicates of the RRS version. Feature importance was quantified as mean decrease in impurity by extracting the values from the attribute ‘feature_importances_’ from the fitted RF model. Similarly, feature importance was also plotted by averaging feature importance of each model built with the triplicates of each RRS version. For the final model, the whole dataset, without splitting, was used to train an RF with 1,000 decision trees with random seed set at 0. A median imputer was also fitted on the whole dataset so that missing features in input protein pairs can be imputed with the median of the corresponding features in the training data. The effect of imputation on the accuracy of the predictor was tested by imputing masked values in the test set, and the effect was found to be negligible. The scikit-learn package from Python was used for model training, feature importance analysis, and ROC and PR statistics^65^.

### Prediction of DMIs in HuRI and comparison to known DMIs

As RRSv4 demonstrated the best performance, the final model was trained using the PRS combined with the RRSv4. To deploy the trained model for DMI scoring, the feature annotation and model inference functions were wrapped into an automated prediction pipeline. The pipeline was then applied on all PPIs found in the HuRI network^16^. For each DMI match, the model outputs a DMI score ranging from 0 to 1 that quantifies the likelihood of the DMI match being functional. A cutoff of 0.7 was used to retain DMI predictions of high accuracy. Only DMI types with human taxonomy annotation (268 in total) were used for generating DMI predictions. We checked if previously annotated ELM instances were also present in the predictions by comparing direct protein pair UniProt IDs and motif and domain residue positions.

### Determination of missense variants overlapping with predicted DMIs

For high-confidence DMI predictions (DMI score ≥0.7), we assessed the overlap with the filtered ClinVar dataset. For each domain and motif (fragment) in the DMI dataset, we identified variants falling within the predicted boundaries of the respective fragment. If a domain consisted of multiple repeats, a variant was considered a match with that domain if it overlapped with any of its repeat matches.

### Cell line culture and maintenance

HEK293 cells were originally obtained from DSMZ (catalog number ACC305) and maintained under standard conditions. Cells were cultured in DMEM (Thermo Fisher Scientific, 11574486), supplemented with 10% FBS (PAN-Biotech, P40-47500) and 1% penicillin– streptomycinPenicillin/Streptomycin(Thermo Fisher Scientific, 15140122). Cultures were kept under 37°C in a 85% humidified atmosphere with 5% CO₂. Routine passaging was performed every 2-3 days using 1mL of 0.05% trypsin (Thermo Fisher Scientific, 25300054), with 0.8 x 10⁶ cells seeded per T25 flask (Sarstedt, 83.3910.002). The purchased cell line was initially expanded, aliquoted and frozen for storage. For quality control, cultures were screened for mycoplasma (Eurofins) contamination once a year using the mycoplasmacheck from Eurofins. No additional authentication of the cell line was performed.

### Plasmid construction

#### Standard controls

The donor vector pcDNA3.1-cmyc-NL-GW (Addgene ID #113446) and control plasmids pcDNA3.1(+), pcDNA3.1-NL-cmyc (Addgene ID #113442), and pcDNA3.1-PA-mCit (Addgene ID #113443) were kindly provided by the Wanker group (Max Delbrück Center for Molecular Medicine, Germany). The acceptor vector pcDNA3.1-mCit-His3C-GW was generated in house.

#### GATEWAY cloning procedure

1. Full-length wild-type human open reading frames (ORFs) cloned in GATEWAY-compatible entry vectors (ORFeome Collaboration, 2016) were maintained as glycerol stock. A sterile swab from each bacterial glycerol stock containing human ORFs was used to inoculate individual wells of 96-well plates, each containing 200 µl of LB medium supplemented with appropriate antibiotics. To minimize evaporation during incubation, the plates were sealed with sterile self-adhesive airpore film (Fisher Scientific, 10642674). Cultures were grown overnight at 37 °C with shaking at 190 rpm.
2. The inoculated ORF cultures were used directly as template for PCR amplification. In a 96-well PCR plate (Starlab, I1402-9700) 4µl of the inoculated culture was used per each 50µl PCR reaction using Phusion High-Fidelity DNA Polymerase (NEB, M0530S), and primers annealing to vector flanking regions (forward: 5′-TTGTAAAACGACGGCCAGTC; reverse: 5′-GCCAGGAAACAGCTATGACC). Thermal cycling was carried out at 98 °C for 10 s, 55 °C for 30 s, and 72 °C for 3 min for 30 cycles.
3. PCR products (6 µl per well) were analyzed on 96-well E-Gel 1% agarose gels containing Ethidium Bromide (Thermo Fisher Scientific, Cat. no. G700801). Each well was loaded with 25 µl of loading buffer (Thermo Fisher Scientific,10482055), and product sizes were assessed using 20 µl of E-Gel 96 High Range DNA marker (Thermo Fisher Scientific, 12352019).
4. For downstream cloning, 1 µl of each amplified PCR product was mixed with 200 ng of the above-mentioned destination vector in a 10 µl Gateway LR recombination reaction using 4× LR Clonase enzyme mix (Thermo Fisher Scientific, 11791020), performed in a 96-well PCR plate and incubated overnight at 25°C to generate the final expression constructs. For increased efficiency 1µl Proteinase K (2µg/µl) was added after the LR reaction and incubated for 10 min at 37°C.
5. The entire 10 µl LR reaction mixture was subsequently transformed into 30 µl of chemically competent *E. coli* DH5α cells (Thermo Fisher Scientific, 18265017) in a 96-well PCR plate. Following transformation, cells were recovered in 80 µl of SOC medium at 37 °C for 1 h without shaking. A total of 70 µl from each transformation was plated onto square 48-well LB agar plates and incubated overnight at 37 °C to allow colony formation.
6. Individual colonies were manually picked and cultured in 96-deep-well plates (Starlab, E2896-2110) containing 2 ml of LB medium supplemented with 100 µg/ml ampicillin (Sigma Aldrich, A9518). Cultures were incubated at 37 °C with shaking at 700 rpm for 24 h in an IncuMixer.
7. Plasmid DNA was extracted using the Plasmid Plus 96 Miniprep kit (Qiagen, 27291) following the manufacturer’s protocol. DNA concentrations were determined using a Nanophotometer, and samples were diluted to a final concentration of 100 ng/µl.
8. For sequence validation, 600 ng of plasmid DNA was subjected to full-length Sanger sequencing using both backbone-specific primers (NanoLuc forward: 5′-GAACGGCAACAAAATTATCGAC; mCitrine forward: 5′-AGCAGAATACGCCCATCG, and reverse: 5′-GGCAACTAGAAGGCACAGTC) and ORF-specific primers (Table S9). All sequence-validated ORFs utilized in this study along with their sequences are listed in Table S10.

#### Site-directed mutagenesis

Primers were manually designed based on the following specifications:

1. For point mutations, primers should be centered on the mutation site with an overlap of 15–20 nucleotides.
2. In cases of deletions, primers should be designed to exclude the deleted region while maintaining similar overlap lengths as stated in step 1.
3. All primers should be 32–36 nucleotides long, with a GC content between 40–60%. Melting temperature (Tm) differences between primer pairs should not exceed 5 °C.
4. The primers should be preferentially designed to start and end with a G or C.
5. The primers should be grouped according to annealing temperature for downstream applications.
6. Site-directed mutagenesis was performed in a 96-well PCR format using 10 ng of plasmid DNA template and primer pairs in 50 µl reactions. Amplification was carried out using Phusion High-Fidelity DNA Polymerase (NEB, M0530S) under the following conditions: initial denaturation at 98 °C for 2 min; 25 cycles of 98 °C for 15 s, annealing (temperature optimized per primer group) for 15 s, and extension at 72 °C for 5 min.
7. To remove template DNA, 1 µl of DpnI (NEB, R0176S) was added directly to each reaction and incubated at 37 °C for 1 h, followed by enzyme inactivation at 65 °C for 20 min.
8. PCR products (6 µl) were confirmed on 96-well E-Gel (Thermo Fisher Scientific, G700801). Each well was supplemented with 25 µl of loading buffer (Thermo Fisher Scientific, 10482055), and size verification was performed using 20 µl of E-Gel 96 High Range DNA marker (Thermo Fisher, 12352019).
9. Chemically competent *E. coli* DH5α cells (30 µl) were transformed with 3 µl of the DpnI-digested PCR products in a 96-well PCR plate format. Cells were recovered in 80 µl of SOC medium at 37 °C for 1 h without shaking. Following recovery, 70 µl of each transformation was plated onto 48-well square agar plates and incubated overnight at 37 °C.
10. Individual colonies were manually selected and grown in 96-deep-well plates (Starlab, E2896-2110) containing 2 mL LB medium, supplemented with 100 µg/ml ampicillin. Cultures were incubated at 37 °C with shaking at 700 rpm for 24 h using an IncuMixer.
11. Plasmid DNA was isolated using the Plasmid Plus 96 Miniprep kit (Qiagen, 27291), quantified using a Nanophotometer, and normalized to 100 ng/µl.

#### Sequencing

1. Sanger sequencing was performed in a 96-well format to validate the full insert. For each sample, 400–600 ng of plasmid DNA was used together with tag- or ORF-specific primers (Table S9) to ensure complete coverage of the open reading frame and engineered mutation sites. Sequencing reactions were set up using two 96-well PCR plates per run: one for forward and one for reverse primers.
2. For each well, 1 µl of the appropriate sequencing primer was added using a multipette and 1 ml combitip. The following primers were used based on the construct type.
3. Subsequently, 6 µl of diluted plasmid DNA was added manually using a multichannel pipette. Plates were sealed with aluminum foil (Sigma-Aldrich, CLS6570-100EA). Each 96-well PCR plate was submitted as an individual sequencing run via the StarSEQ online submission system.
4. Upon return of sequencing data, reads were processed using an in-house Sanger sequencing analysis pipeline. Sequences of inserts with mutations are available in Table S10.

### BRET assay

#### Transfection

HEK293 cells were cultured in high-glucose (4.5 g/l) DMEM (Thermo Fisher Scientific, 11574486), supplemented with 10% FBS (PAN-Biotech, P40-47500) and 1% Penicillin/Streptomycin (Thermo Fisher Scientific, 15140122). Cells were maintained at 37 °C with 5% CO₂ and 85% relative humidity and subcultured every 2–3 days. For BRET assays, reverse transfection was performed using Lipofectamine 2000 (Thermo Fisher Scientific, 11668019) in Opti-MEM (Fisher Scientific, 12559099) according to the manufacturer’s protocol. Cells were seeded in 96-well white flat-bottom plates (VWR, 392-0229) at a density of 4.0 × 10⁴ cells per well in phenol red-free, high-glucose DMEM (Thermo Fisher Scientific, 31053028) supplemented with 5% FBS (PAN-Biotech, P40-47500) and 1% Pen/Strep. Transfections were carried out with a total of 200 ng plasmid DNA per well. When the DNA amount of expression plasmids was below 200 ng, pcDNA3.1(+) was added as carrier DNA to reach the total amount. Prior to transfection, plate layouts were designed including appropriate controls: NL-stop + pcDNA, mCit-stop + pcDNA, pcDNA-only wells, and wells with untransfected cells. Control DNAs (PA-mCit-Stop, NL-Stop, pcDNA3.1) were prepared in PCR strips (Starlab, I1402-3700) for efficient pipetting. NL-constructs were further diluted to 4 ng/µl. Cells were harvested, trypsinized, counted, centrifuged, and adjusted to a final concentration of 2.67 × 10⁵ cells/ml in phenol red-free DMEM and 150µl of this cell suspension was pipet into each well. Plates with the transfected cells were incubated at 37 °C with 5% CO₂ and 85% relative humidity for 48 hours prior to measurement.

#### Measurement

All measurements were performed using an Infinite M200 Pro microplate reader (Tecan). First, 100 µl of the medium were aspirated from each well. mCitrine fluorescence (FL) was measured in intact cells using excitation/emission wavelengths of 513/548 nm, a gain setting of 100, and 4 point measurements per well, which were averaged to represent the FL value for this well. Following fluorescence measurement, coelenterazine-h (PJK Biotech GmbH, 102181) was added to each well at a final concentration of 5 µM. The plate was briefly shaken for 15 s and incubated for 15 min inside the plate reader at 37 °C. After incubation, total luminescence (totLu) was measured per well, followed by short-wavelength and long-wavelength luminescence using the BLUE1 (370–480 nm) and GREEN1 (520–570 nm) dual filter setting, respectively, with an integration time of 1000 ms. Transfections with NL-stop at 2 ng in one well were used to determine the bleedthrough. Transfections with pcDNA3.1 at 200 ng in one well were used to determine background totLu and FL signals.

#### BRET titrations

For each protein pair the following six different donor:acceptor DNA ratios were used for transfections: 2:12.5 ng, 2:25 ng, 2:50 ng, 2:100 ng, 1:100 ng, and 1:200 ng. Each well contained 200 ng of DNA in total, with pcDNA3.1(+) added to the well to reach this amount. DNA mixtures for each protein pair were first prepared as master stocks in a 96-well PCR plate for three technical replicates. From this master plate, the DNA mixtures were distributed into three 96-well white flat-bottom plates (VWR, 392-0229). Transfection and subsequent measurements were conducted as described above.

### Quantification of BRET signals

Average background FL and totLu signals (see above) from one biological replicate were subtracted from each measured FL and totLu value, respectively. If no background FL and totLu values were available for a given biological replicate, average values across all experiments were used instead (299 for FL, 7279 for totLu). The average bleedthrough (see above) from an experiment was subtracted from the BRET signals measured for each well. If no bleedthrough measurements were available for a given biological replicate, the average as determined across all experiments was used instead (0.297). After background correction all negative FL, totLu, and BRET values were set to zero. For every protein pair six titration points per technical replicate were measured (see above). Titration points were excluded, if totLu < 15000. Titration points were fitted, if at least five titration points for a given technical replicate passed this cutoff and at least three of these titration points had FL values ≥ 300. These filtering criteria were put in place to ensure that only titration points from measurements with sufficient donor (totLu) and acceptor (FL) expression levels were used to derive a fit.

The leastsq function from the scipy.optimize python package was used to fit the remaining titration points per replicate to a simple 1:1 binding model with BRET = ((A/D) * BRETmax)/(BRET50 + (A/D)) where A = acceptor protein levels (FL) and D = donor protein levels (totLu) with the goal to obtain estimates for the BRETmax and BRET50 parameters^66^. BRET50 corresponds to the A/D ratio at which half of the BRETmax is reached. BRETmax is the estimated maximal BRET that can be reached for the given protein pair upon saturation. Standard errors of the BRET50 and BRETmax estimates were obtained from the variance-covariance matrix, calculated by multiplying the fractional covariance matrix (output by leastsq function) by the residual variance. A fit was deemed valid, if the estimated BRET50 was smaller than the max expression ratio (FL/totLu) that was measured for the given titration curve and if the max measured BRET for that titration curve was ≥ 0.05. These criteria were put in place to identify accurate fits, which can be obtained if saturation is (close to being) observed (that is not the case if the BRET50 is > the max expression ratio) and if at least one BRET value indicates binding (BRET > 0.05).

If for a given protein pair at least three fits are valid, then BRET50 values can be averaged to quantify interaction strength of that pair. If for a given wildtype protein pair and corresponding mutant pair at least three valid fits are available for each pair from the same biological replicate, then student’s T-test was applied on the wildtype and mutant BRET50 values to quantify significant changes in interaction strength between wildtype and mutant. If less than three valid fits were available for wildtype or mutant protein pair for a given biological replicate, then differences in interaction strength could not be quantified using BRET50. This was the case where mutant or/and wildtype protein pairs did not reach saturation or where the mutant protein was very disruptive to protein binding. In these cases and as long as at least three technical replicates of titration curves each from wildtype and mutant protein pair were valid (but not the fit) we determined differences in the maxBRET as follows. The highest expression ratio (FL/totLu, hereafter referred to as max_expr_ratio) that was obtained for wildtype or mutant pair across the three technical replicates of the titration curves was determined. Consequently, within a sliding window of size max_expr_ratio/4 and a step size of the sliding window of max_expr_ratio/20 the highest expression ratio window was identified that contained at least three BRET measurements each from wildtype and mutant pair. These BRET values were taken to determine significant differences between wildtype and mutant protein pair using the student’s T-test. The rationale behind this approach is to compare BRET values between wildtype and mutant protein pair that were obtained for similar expression ratios excluding the possibility that differences in BRET were obtained from differences in expression levels. We furthermore assume that the tested mutations did not significantly alter the overall conformations of the proteins (due to testing interface mutations) why differences in BRET between wildtype and mutant protein pair most likely result from differences in interaction strength. If for a given wildtype and mutant protein pair BRET50 comparisons could not be conducted (see above) then differences in maxBRET were used instead.

### Structural modeling of experimentally tested PPIs

The potential DMIs of experimentally tested PPIs were subject to structural modeling using a local installation of AlphaFold Multimer version 2.3.2^19^. Lee et al. 2024^20^ demonstrated that including the flanking residues around motifs in AlphaFold predictions could improve the accuracy of predicted models. Therefore, to ensure the most optimal configuration, we extended the motif matches by the length of the motif match at the N- and C-terminus of the motif match unless the protein termini were reached before in which case extensions until the protein terminus were used. For the domains, we took the positions of their HMM matches and extended them to include 10 more residues at the N- and C-terminus to account for the often inaccurate domain boundaries predicted by HMMs. In cases where tandem repeats are involved, the start of the first tandem repeat and the end of the last tandem repeat were used as the domain match and similar extensions of 10 residues were applied on them. The extended motif and domain sequences were paired and submitted to AlphaFold Multimer for structural modeling.

For the negative control PPIs that are without any DMI match, the fragmentation approach described in Lee et al. 2024^20^ was applied to predict the potential mode of binding. In brief, the predicted monomeric structures of the individual proteins in the PPIs were downloaded from AlphaFold database^55^, and they were manually inspected for folded and disordered regions. Following that, the disordered regions were fragmented into 20 residue-long fragments with a sliding window of 10 residues to generate overlapping fragments of disordered regions. Finally, for each PPI, we paired the fragmented disordered regions of one protein with the folded regions of its interactors and vice versa as well as ordered with ordered regions for AlphaFold Multimer structural modeling. To evaluate the modeled structures, we applied a cutoff of 0.7 model confidence and structures surviving this cutoff were manually inspected prior to final approval. The manual inspection of modeled structures involved checking if overlapping fragments were repeatedly modeled at the same binding site, surveying literature for existing or potential binding pockets at the modeling binding sites, and the plausibility of the interchain interactions modeled at the binding sites.

The local installation of AlphaFold Multimer version 2.3.2 was run using the following parameters:

--model_preset=multimer

--max_template_date=2020-05-14

--db_preset=full_dbs

--use_gpu_relax=False

For every AlphaFold run, five models were predicted with single seed per model by setting the following parameter:

--num_multimer_predictions_per_model=1

The databases queried during AlphaFold predictions were specified following the instructions from the github page of AlphaFold (https://github.com/deepmind/alphafold#running-alphafold):

--bfd_database_path=alphafold_v232_databases/bfd/bfd_metaclust_clu_complete_id30_c90_ final_seq.sorted_opt

--mgnify_database_path=alphafold_v232_databases/mgnify/mgy_clusters_2022_05.fa

--obsolete_pdbs_path=alphafold_v232_databases/pdb_mmcif/obsolete.dat

--pdb_seqres_database_path=alphafold_v232_databases/pdb_seqres/pdb_seqres.txt

--template_mmcif_dir=alphafold_v232_databases/pdb_mmcif/mmcif_files

--uniprot_database_path=alphafold_v232_databases/uniprot/uniprot.fasta

--uniref30_database_path=alphafold_v232_databases/uniref30/UniRef30_2021_03

--uniref90_database_path=alphafold_v232_databases/uniref90/uniref90.fasta

For the fragmentation approach, the multiple sequence alignments (MSAs) of a given protein fragment can be reused in subsequent runs where the same fragment is involved. The MSAs were first moved to the prediction output folder and the following parameter was added to enable the reuse of MSAs.

--use_precomputed_msas=True

For efficient computing, we segregated the MSA generation part by using only the CPUs and the model fitting part using the GPUs.

### Mutation design to validate predicted DMIs

To experimentally validate predicted DMIs, we designed mutations based on the structural models generated using AlphaFold (see above). These models were analysed with the PyMOL tool^67^. Guided by the predicted interfaces, we manually selected mutations to disrupt domain or motif interfaces. Mutation types included single amino acid substitutions at key interface residues as well as deletions of entire motifs. Mutations were introduced into the coding sequences using site-directed mutagenesis as described above. All mutant constructs were tested for expression using totLu or FL measurements (see above), and only those that showed expression levels comparable to their wild-type counterparts were included in downstream validation assays (see BRET titration assay section). Constructs with insufficient expression were excluded from further experiments. All structural model graphics were produced using ChimeraX^68^.

### Generation of multiple sequence alignments

MSA sequences were first retrieved from ProViz^69^ using their UniProt accession. We aimed to construct a representative group of 12 vertebrates and in cases of missing sequences, we attempted to retrieve them by homology search using BLAST. All sequences were realigned in Jalview^70^ using the T-Coffee default parameters^71^ and visualised using Clustal colors.

### Structural modelling of predicted DMIs

From all predicted DMIs that met the 0.7 DMI score cutoff we selected 10,228 predicted DMIs that contained at least one pathogenic or uncertain variant in either fragment. From those we retained 5,463 predicted DMIs that did not require post-translational modifications on motif residues for partner recognition since they cannot be modelled by AlphaFold v2.3, and excluded all DMI types of the ELM category MOD. When protein pairs contained multiple matches for the same DMI type because of repeated domains, each domain was modelled individually. This resulted in 7,682 high-scored DMIs for structural modelling. The sequence boundaries of domains (defined in the DMI predictor based on HMM matches) were expanded using pLDDT values of human AFDB structures (July 15th 2024) to include missing secondary structure elements of the folds. The pLDDT values of the first and last 10 residues of an HMM match were averaged and if the mean was equal or above a pLDDT threshold of 80 the domain boundaries were extended by one position and the mean recalculated until the threshold was not met anymore. For motif sequences we took the sequence REGEX match and added four residues upstream and five residues downstream when possible. The fragment sequence of each DMI was subjected to AlphaFold modelling using the same local installation and parameters as mentioned above.

### Classification of residues in structural models

We took the PDB files from the relaxed best score model of each AF DMI prediction and also saved both chains in individual PDB files. We took the three PDB files to calculate total surface accessibility through DSSP^72,73^. pLDDT and accessibility values were retrieved from model .cif files and DSSP table files, respectively. Relative surface accessibility (RSA) was calculated by dividing their total accessibility by the theoretical maximal accessibility using the residue normalisation scale from^74^. Delta RSA was calculated by subtracting the residue RSA of the unbound apo structures (single chain files) from the bound holo structure (complex file).

Structural classification of residues within the models was done by the following criteria: residues in either domain or in core motif chains (residues that correspond to the regular expression match) with pLDDT values greater or equal 80 and a delta RSA equal or smaller than -0.05 were considered interface. Motif residues not following these criteria as well as flanking residues were considered not-interfacial. Domain residues not following the criteria were further classified into buried if their RSA was ≤ 0.25 on the holo structure and surface if they had a RSA > 0.25^75^.

### AlphaMissense predictions

The AlphaMissense_aa_substitutions table was retrieved from https://zenodo.org/records/8208688 (July 29th 2024)^22^. The AlphaMissense amino acid change pathogenicity scores, as well as their assigned discrete categories, were mapped to ClinVar variants using protein UniProt ID, residue position number and amino acid information.

### Retrieval of disease annotations and DMI-based network

We focused on proteins from DMI models containing uncertain variants at interfacial residues that AlphaMissense classified as benign. Association of these proteins to a direct genetic disease from OrphaNet (https://www.orphadata.com/orphadata-api/) (February 21st 2025)^76^ was done using protein gene names. In order to group the different genetic diseases into larger categories we employ Disease Ontology (DO)^77^ parent terms using the “rols” R library^78^. Using the total count of occurrences of parent terms, the ones shared among mapped diseases, we kept one parent term per gene and took 6 of the top 20 terms that represented different disease etiologies. Using the DMI PPI pair information we generated an interaction network using Cytoscape^79^. The PPI network was visualised using a Compound Spring Embedder (CoSE) layout with manual adjustment to improve label visualization.

---

**Extended data figure 1.**
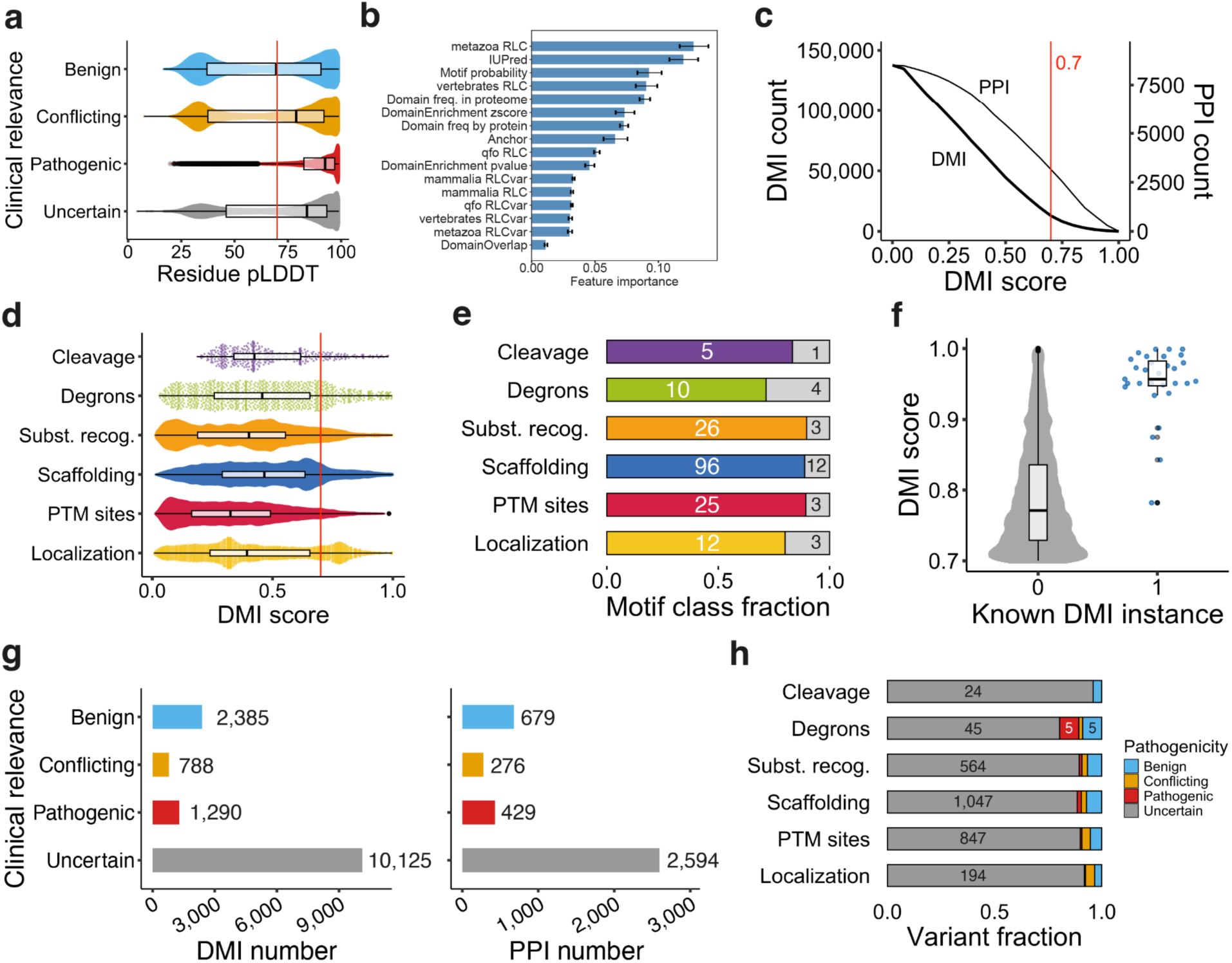
Prediction of DMIs and overlap with variants. **a.** pLDDT distribution of variants by clinical significance. The red line indicates a pLDDT cutoff of 70 used to distinguish disordered from ordered residues. **b.** Feature importance of the Random Forest model trained using the PRS and RRS4 as quantified using mean decrease in impurity. **c.** Number of predicted DMIs and PPIs with at least one predicted DMI for increasing DMI predictor score cutoffs. The red line indicates the used DMI score cutoff of 0.7 corresponding to 12,957 predicted DMIs in 3,188 PPIs. **d.** Distribution of DMI predictor scores of predicted DMIs by motif category. The red line indicates the used DMI score cutoff of 0.7. **e.** Number of DMI types by motif category. Colored bars correspond to DMI types with at least one predicted DMI that passed the 0.7 cutoff. **f**. Distribution of DMI scores for predicted DMIs that do (1) or do not (0) correspond to annotated DMIs in the ELM resource. **g.** Number of predicted DMIs (left) and corresponding PPIs (right) that overlapped with at least one variant of indicated clinical significance. **h.** Fraction and number of variants that overlapped with at least one predicted DMI by motif category.

**Extended data figure 2.**
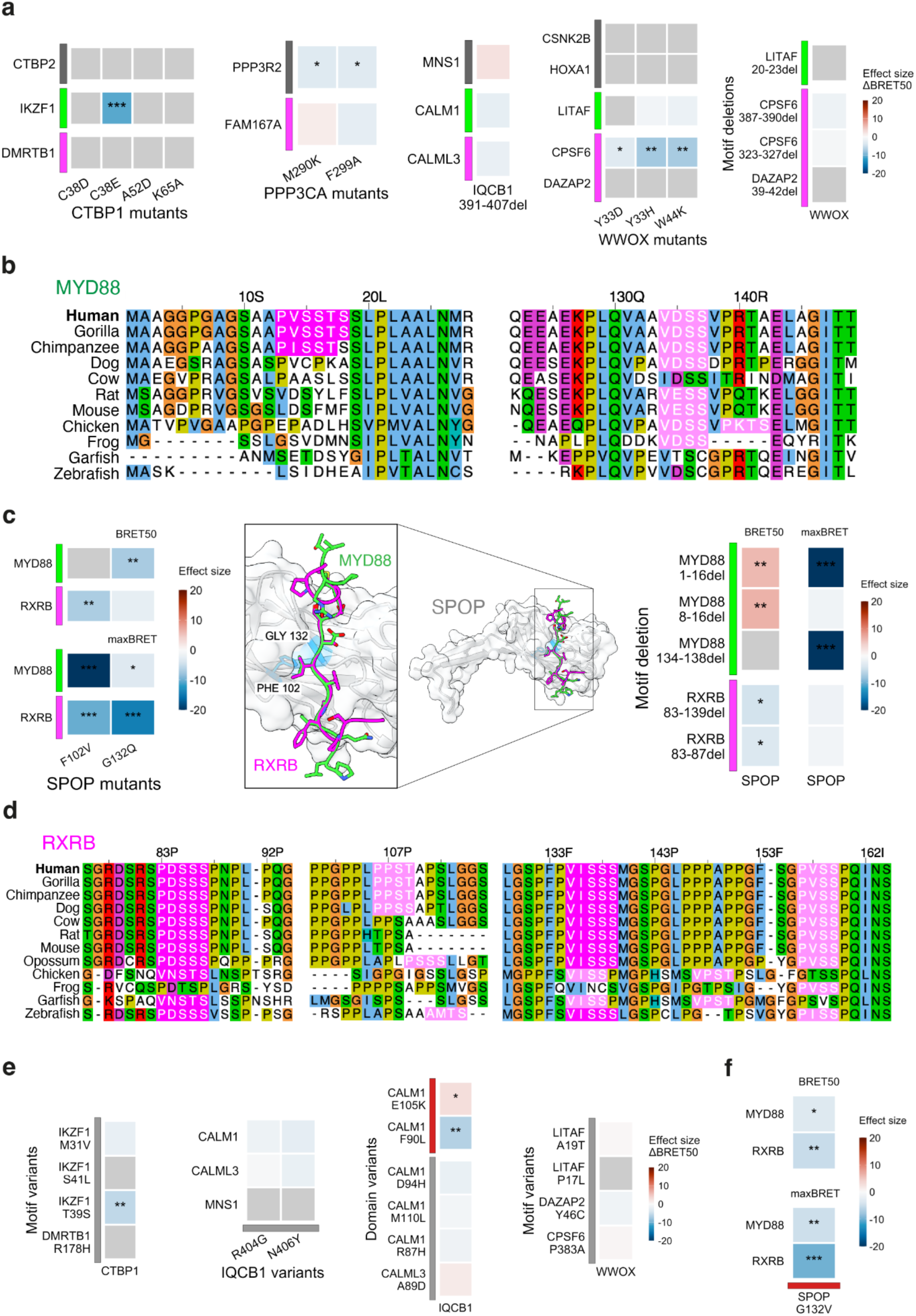
Experimental validation of predicted DMIs and effects of variants on PPIs. **a**. Heatmap of effect sizes based on T-test of ΔBRET50 estimates obtained from fitting wildtype and mutant BRET titration curves to validate predicted DMIs. Grey=no BRET50 estimates available because titration curves could not be fitted. *** p-value < 0.001, ** p-value < 0.01, * p-value < 0.05, three technical replicates each. Effect size values capped at 20 and -20. Color bars on y-axis: grey=other interface, green=known DMI, magenta=predicted DMI. **b.** MYD88 multiple sequence alignment of regions carrying the predicted DMI below the 0.7 cutoff (highlighted in magenta) and the known DMI (highlighted in light pink) that did not match the regular expression for the SPOP degron. **c.** Heatmaps as in **a** but for both estimated ΔBRET50 and measured ΔmaxBRET between wildtype and mutant SPOP interactions with RXRB or MYD88 partner proteins to assess predicted and confirm known motifs in RXRB and MYD88, respectively. Superimposed structural models of the MATH domain of SPOP (1-175) with MYD88 (128-143) and RXRB (130-144) SPOP degron motifs. MATH domain residues mutated to validate the motif-binding pocket are highlighted in blue. **d.** RXRB multiple sequence alignment of regions carrying predicted motifs that match the SPOP degron regular expression (magenta) and putative motifs that do not match the regular expression (light pink). **e-f.** Heatmaps as in **a** and **c** but showing effects of ClinVar variants on PPIs. Color bars on y- or x-axis: grey=uncertain variants, red=pathogenic variants.

**Extended data Figure 3.**
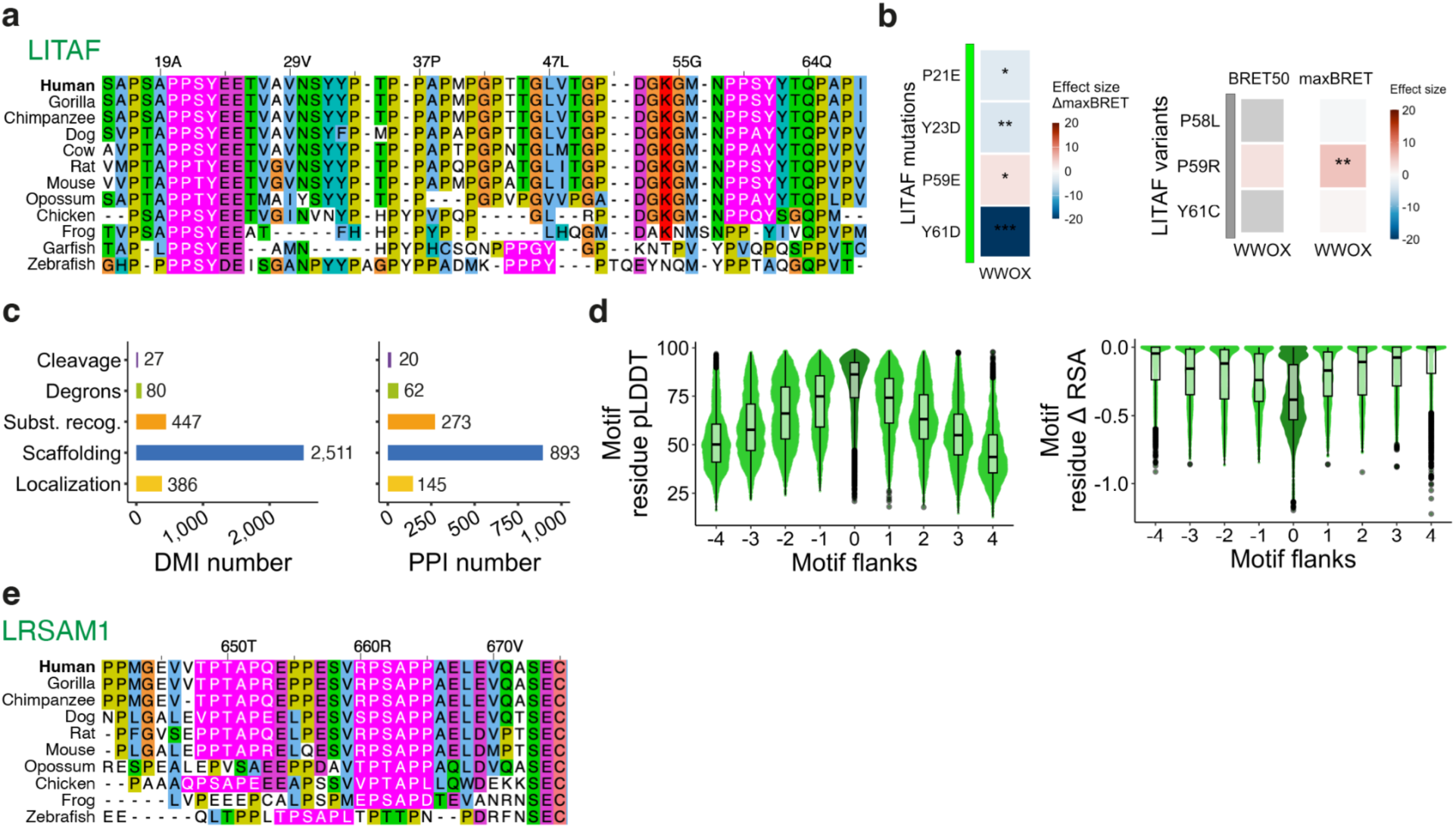
Multiple sequence alignments and structural modelling of predicted DMIs. **a.** LITAF multiple sequence alignment showing two conserved PPxY motifs highlighted in magenta. **b.** Heatmap of effect sizes based on T-test of ΔBRET50 estimates and ΔmaxBRET measurements obtained between wildtype and mutants to probe disruption of known motifs and uncertain variants in LITAF falling within the motif when binding to WWOX. *** p-value < 0.001, ** p-value < 0.01, * p-value < 0.05, three technical replicates each. Effect size values capped at 20 and -20. Color bars on y-axis: green=known interface, grey=uncertain variants. Grey fields in heatmap indicate protein pairs for which no BRET50 estimates could be obtained. **c.** Number of predicted DMIs (left) and corresponding PPIs (right) per motif category with confident structural models (model confidence > 0.6). **d.** pLDDT (left) and ΔRSA values of motif residues from confident structural models of predicted DMIs. Core motif residues (those that match the regular expression) are averaged as position 0. Upstream and downstream residues from the core motif are labeled by negative and positive residue positions, respectively. **e.** LRSAM1 multiple sequence alignment showing two conserved PTAP motifs (pink).

